# Identification of sodium/myo-inositol transporter 1 as a major determinant of arterial contractility

**DOI:** 10.64898/2025.12.12.694071

**Authors:** Elizabeth A Forrester, Christopher J Garland, Vincenzo Barrese, Anthony P Albert, Iain A Greenwood

## Abstract

**Background:** As the sodium/ myoinositol transporter (SMIT1) is a positive regulator of Kv7.4/7.5 channels in arterial smooth muscle we postulated that altering SMIT1 expression could have a major impact upon vascular reactivity. Consequently, this study aimed to characterise the effects of changes of SMIT1 membrane abundance on vascular tone and the molecular mechanisms involved.

**Methods:** Isometric tension recording on 2^nd^ order mesenteric arteries and left anterior coronary arteries from male and female Wistar Han rats. Whole artery membrane potential recording. Single cell and whole artery antibody-based imaging. Morpholino based protein knockdown of SMIT1.

**Results:** Morpholino-mediated knockdown of SMIT1 enhanced U46619- and methoxamine-mediated contractions of mesenteric artery whilst impairing relaxations to the Kv7 activator ML213, isoprenaline and CGRP. Conversely, augmenting SMIT1 membrane abundance by raising external osmolarity with 150 mM raffinose, impaired receptor-mediated contractions of mesenteric and coronary arteries, augmented relaxations to ML213 and adenosine as well as producing membrane potential hyperpolarisation. Proximity ligation assays revealed that raffinose incubation increased the association of SMIT-Kv7.4/7.5 as well as Kv7.4 and Gβγ subunits. The SGK1 inhibitor EMD638683 prevented the raffinose-induced increase in SMIT1 and the anti-contractile effect.

**Conclusion:** Toggling the membrane abundance of SMIT1 had a dramatic effect on the arterial response to vasonstrictors and vasodilators mediated by greater coordination of Kv7 channels and Gβγ subunits. This work identified SMT1-Kv7 channel complexes and SGK1 regulation as key modulators of arterial responsiveness.

## 1. Introduction

The sodium/myoinositol transporter (SMIT1) encoded by SCL5a3 provides substrate for the synthesis of the minor phospholipid phosphatidylinositol bisphosphate (PIP_2_), which has important role in cellular signalling and is a crucial co-factor for many ion channels^1^. SMIT1 has also been identified to alter the ion selectivity, gating, and pharmacology of KCNQ-encoded voltage gated potassium channels (Kv7.1-7.5) independent of inositol transport through a direct physical interaction with the pore component ^2–7^. These channel-transporter couples (‘chansporters’) have been implicated as important regulators of neuronal activity at rest and in response to increase external osmolarity ^8–10^. SCL5a3 gene expression is enhanced by a tonicity-responsive enhancer binding protein (TonEBP, ^11^) leading to increased expression of SMIT1. In addition, SMIT1 protein levels in the cell membrane are influenced by ubiquitin ligases that are down-regulated by hypertonically-regulated serum-glucocortoid sensitive kinase (SGK) 1 ^10^, which offers a rapid mechanism to alter SMIT levels.

In arterial smooth muscle activating Kv7 channels prohibits arterial contraction and contributes to receptor-mediated vasodilatation ^12–14^. SMIT1 associates with Kv7.4 and Kv7.5 in smooth renal artery muscle cells and overnight incubation in hypertonic medium increased SMIT1 expression markedly, augmented Kv7 channel currents and reduced vasoconstrictor responses^6^. The juxtaposition of SMIT regulation by hypertonicity and a role as a large auxiliary channel for Kv7 channels offers an effective mechanism to transduce changes in external environmental status to arterial responses. We hypothesised that altering SMIT1 membrane abundance would modulate responses to receptor-mediated dilators and offered a mechanism to fine tune arterial contractility. Our study provides clear evidence that SMIT1 is a key regulator of vascular responses that has connotations for vascular disease development.

## 2. Methods

Experiments were performed on second-order mesenteric and cardiac septal resistance arteries from male and female rats aged 11–14 weeks and weighing 175–300 g. Animals were maintained under an institutional site licence and sacrificed by a Schedule 1 method (cervical dislocation) in accordance with the UK Animal (Scientific Procedures) Act 1986; therefore, no approval from a local or university ethics review board was required. This investigation conforms to Directive 2010/63/EU of the European Parliament on the protection of animals used for scientific purposes. All methods for isometric tension and membrane potential recording, morpholino-based protein knockdown, immunohistochemistry and PCR are described fully in the supplemental materials.

### 2.1. Data and statistical analysis

All values from functional experiments were expressed as mean ± standard error of the mean (SEM) with no less than 5 individual data points, each representing a biological repeat. Measurements of total cell fluorescence during immunocytochemistry involved five biological repeats with a minimum of five cells recorded per sample. For functional experiments, cumulative concentration effect curves were produced. The tone of the artery was recorded after each subsequent addition of a given pharmacological agent and the values were formulated as a percentage of the maximum contraction. Using GraphPad Prism (RRID:SCR_002798, Version 10.0.0) a transformed data set of mean values was generated using X = Log(X), to reduce representative skew. A four parametric linear regression analysis was then performed to produce a concentration effect curve on a log(x) graph with the standard error of mean (SEM). A two-way ANOVA was performed followed by a post-hoc Bonferonni test for multiple comparisons or one-way Anova were used to determine statistical significance between groups, according to the different experiments. Significance values were represented as * = P<0.05, ** = P<0.01, *** = P<0.001 and **** = P<0.0001. All data sets subject to statistical analysis contained at least 5 animals per group, where N = number of independent values.

## 3. Results

### 3.1. Downregulating SMIT1 levels increases arterial contractility and impairs the relaxation responses to CGRP and isoprenaline

Quantitative PCR was undertaken to ascertain the relevant abundance of SCL5a3 and SCL5a11 that encode for SMIT1 and SMIT2. Figure 1A shows that renal, mesenteric and coronary arteries express SCL5a3 almost exclusively with negligible contribution from SCL5a11. Whole artery immunohistochemistry showed robust expression of SMIT1 in the smooth muscle (MYH11 positive, 1B) and endothelial PECAM/CD31 positive, 1C) layers. To investigate the functional impact of SMIT1 in mesenteric arteries we used morpholinos blocking the translation of SMIT1 shown by Barrese et al (2020) ^6^ to reduce SMIT1 expression in single cells by roughly 40%. Transfection with SMIT1-specific morpholino reduced SMIT1-positive staining in the smooth muscle and endothelial layers of mesenteric arteries (Fig 1B, C). Figure 1D-E shows that responses to methoxamine and U46619 were enhanced considerably by knockdown of SMIT1 whereas contractions to 60 mM KCl were not affected (Fig 1F). In addition, relaxations to isoprenaline and CGRP were impaired by SMIT1 knockdown (Fig 1G-H) and relaxations produced by 1µM ML213 (Kv7.2-7.5 activator^15–17^) were reduced from 84.55 ± 3.28 % to 27.02 ± 10.38 % (N=5, Fig 1I). These data highlight that reducing SMIT1 impacts on vascular reactivity.

**Figure 1:**
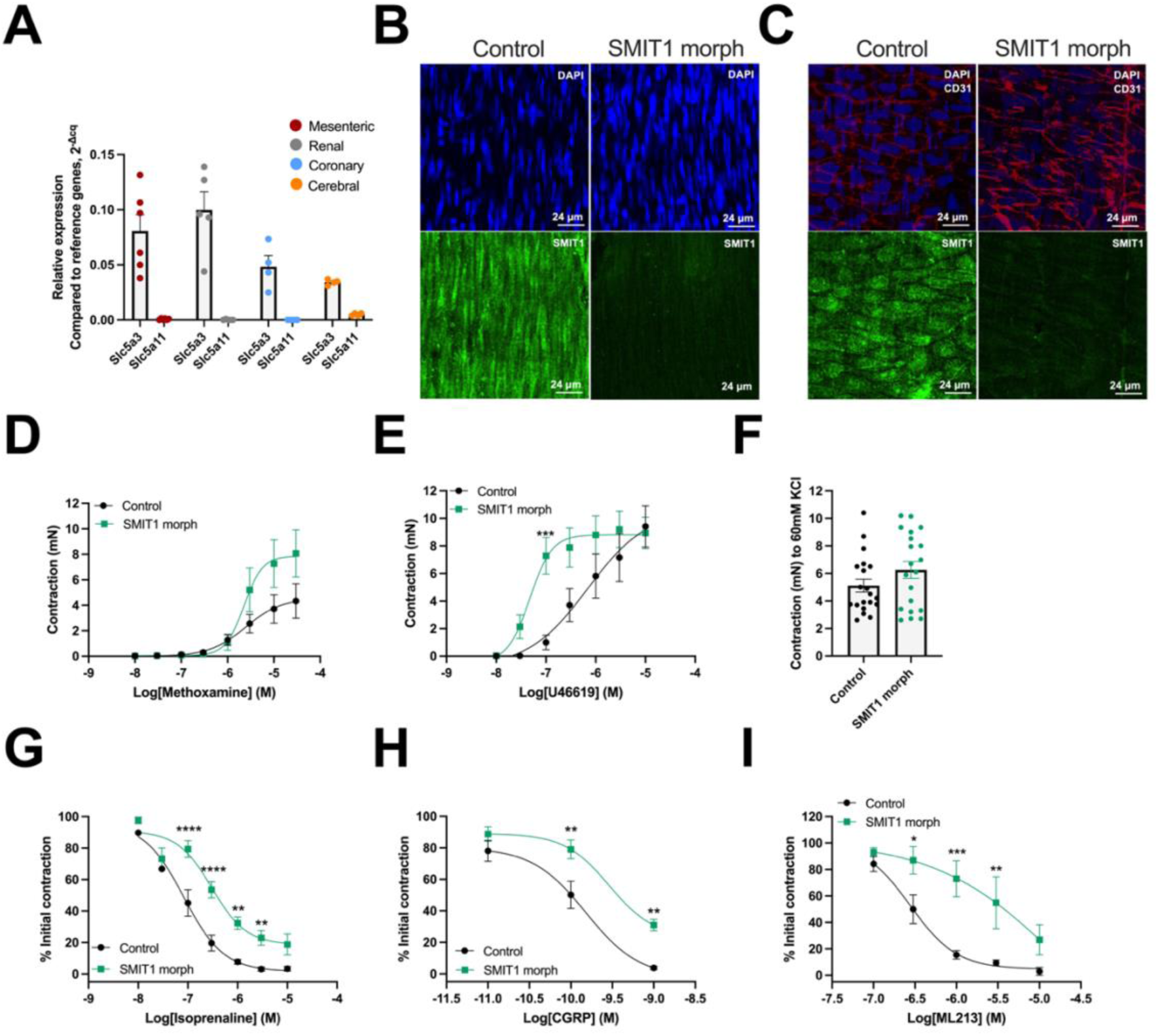
Arterial expression of SMIT1 and the effect of SMIT1 knock-down on arterial contractile and relaxation responses. The mean transcript abundance of Slc5a3 and Slc5a11 in mesenteric (red), renal (grey), coronary (blue) and cerebral (orange) arteries from male rats (A). The expression of SMIT1 protein (green) in smooth muscle cell layers (B) and endothelial cell layers (C) of mesenteric arteries transfected with scrambled control morpholino and SMIT1 targeted morpholino. The endothelial cells are stained with CD31, shown in red, with nuclear DAPI stain in blue. The mean raw concentration-dependent contraction (mN) to methoxamine (D) and U46619 (E) in mesenteric arteries incubated with either a scrambled control morpholino (black) or a SMIT1 targeted morpholino (green)(N=8). The contraction to 60mM KCL in mesenteric arteries incubated with either a scrambled control morpholino (black) or a SMIT1-targeting morpholino (green, F, N=20). The mean concentration-dependent relaxation (mN) to Isoprenaline (1 nM – 1 µM) in scrambled control morpholino (black) and SMIT1 targeted morpholino (green) mesenteric arteries pre-contracted with 300 nM U46619 (G, N=6). The mean concentration-dependent relaxation (mN) to CGRP (H, 10 pM – 1 nM) and ML213 (I, 10 nM – 10 µM) in scrambled control morpholino (black) and SMIT1 targeted morpholino (green) mesenteric arteries precontracted with 300 nM U46619 (N=7). All data points are shown as mean +-SEM denoted by error bars, and statistical values are determined by 2-way ANOVA with multiple comparisons are calculated using post hoc Bonferroni test where *=P<0.05, **=P<0.01, ***=P<0.001, ****=P<0.0001.

### 3.2. Acute incubation of hypertonic raffinose enhances arterial relaxation responses

The expression of SMIT1 is upregulated by a hypertonicity activated promotor element (TonEBP, ^11^). Raffinose (150 mM) is a non-metabolizable sugar that increased osmolarity from 271 mOsm to 480 mOSm. Barrese et al., (2020)^6^ showed that overnight incubation of renal arteries in 150 mM raffinose increased SMIT1 fluorescence intensity in renal artery smooth muscle cells and markedly impaired methoxamine-induced contractions compared to control PSS. We ascertained whether changes in arterial responsiveness could be elicited by shorter hypertonic shocks. Incubation in raffinose containing PSS increased the total SMIT1 expression and the presence of SMIT in the cell membrane (Fig 2A-C). Figure 2D-E shows that incubation in 150 mM raffinose for 10 mins to 6 hours followed by wash out into normal PSS inhibited the contractile response to methoxamine. Incubation in raffinose for 4 hours also significantly attenuated vasoconstrictor responses to U46619 in both mesenteric (Fig 2F) and coronary arteries (Fig 2G). Raffinose treatment also reduced contractions elicited by low concentrations of KCl whereas responses to KCl concentrations above 40 mM were not affected (Fig 2H). Raffinose-induced attenuation of methoxamine was was not dependent upon the presence of a functional endothelium (S1). However, the anti-contractile effect of raffionse was prevented by knockdown of SMIT1 through translation blocking morpholinos (Fig 2I), application of pan-Kv7 blocker XE991 and the Gβγ inhibitor gallein (50µM) in mesenteric (Fig 2J) and coronary (Fig 2K) arteries. These data reveal a rapid anti-contractile effect of raffinose due to an enhanced Kv7 mediated potassium flux driven by increased SMIT1 expression.

**Figure 2:**
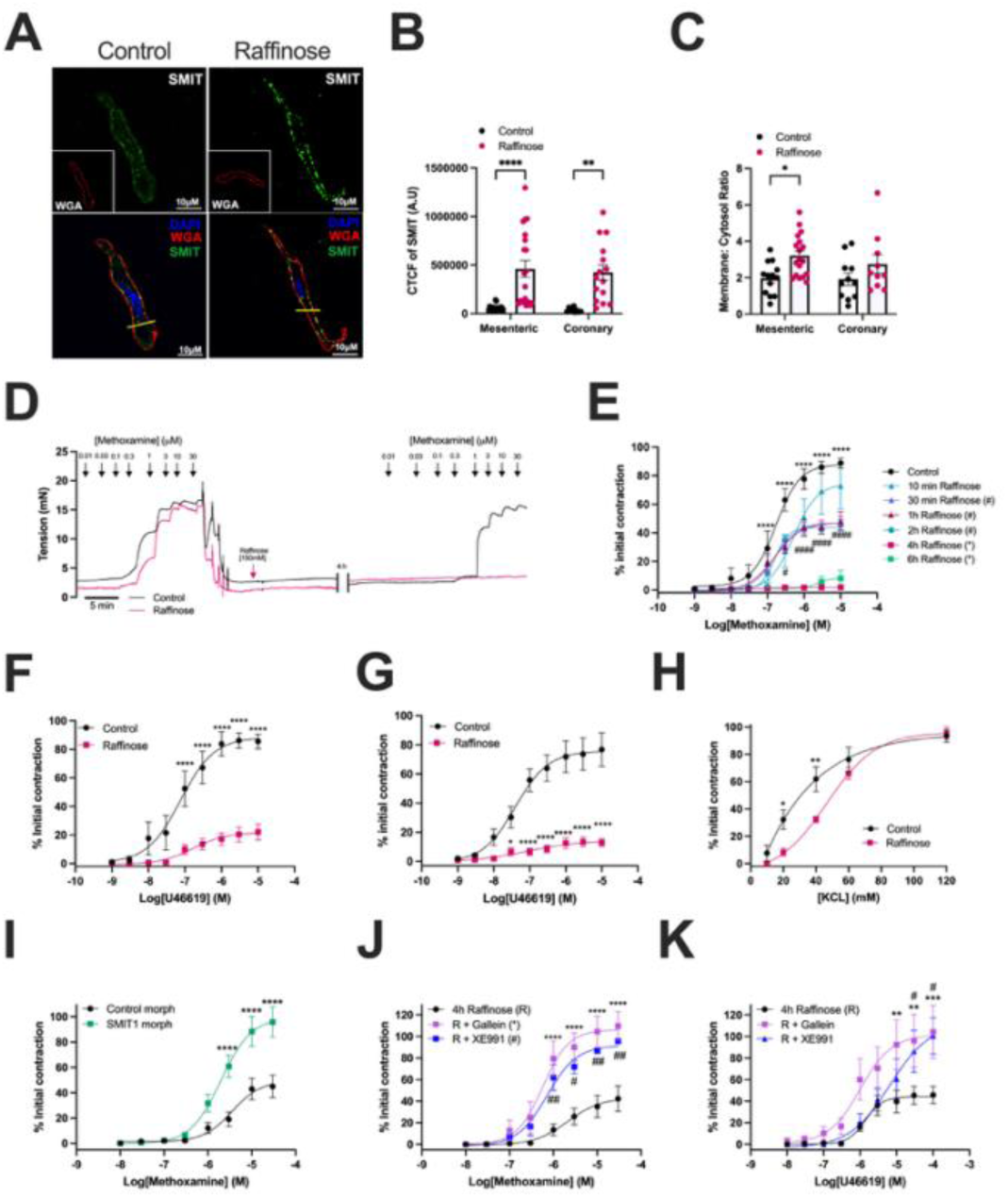
The effect of incubating in hypertonic raffinose on arterial contractile responses. A representative image of male vascular smooth muscle cells stained with SMIT1 (green), membrane marker, wheat germ agglutinin (WGA, red) and nucleus stain with DAPI in control (A, left) and after 4 hours 150 mM raffinose (A, right). The mean corrected total cell fluorescence (CTCF) intensity of SMIT1 (B) and mean membrane cytosol ratio of SMIT1 (C) in mesenteric (N=3, n=15) and coronary (N=3, n=10) arteries incubating in control PSS (black) or 150mM raffinose PSS (pink) for 4 hours. A representative trace of the response to methoxamine (10 nM – 30 µM) in vessels before and after incubating for 4 hours in control PSS (black) and 150mM raffinose (pink, D). The concentration-dependent contraction to methoxamine after incubating in either control PSS (black) or 150mM raffinose for 10 minutes (turquoise), 30 minutes (blue), 1 hour (red), 2 hours (teal), 4 hours (pink) and 6 hours (green) compared to control (black) (N=5-9) shown in panel E. The mean data of the U46619-induced contraction in mesenteric arteries (F) and coronary arteries (G) with (pink, EC50 = 308.5 nM) and without (black, EC50 = 27.33 nM) incubating in raffinose PSS (N=8). The mean concentration dependent response to KCL in the presence (pink) and absence (black) of incubating with 150mM raffinose. The mean concentration-dependent contraction (mN) to methoxamine (10 nM – 10 uM) in scrambled control morpholino (black, EC50 = 3.56 µM) or a SMIT1 targeted morpholino (green, EC50 =1.97 µM) mesenteric arteries incubated in 150 mM raffinose (4 hours) is shown in panel I (N=9). The mean contraction to methoxamine in the presence of 150 mM raffinose (black) with either XE991 (blue) or Gallein (purple) in mesenteric (J) and coronary arteries (K) (N=8).All data points are shown as mean +-SEM denoted by error bars, and statistical values are determined by 2-way ANOVA with multiple comparisons are calculated using post hoc Bonferroni test where *=P<0.05, **=P<0.01, ***=P<0.001, ****=P<0.0001.

As SMIT1 knockdown impaired responses to Gs linked receptors we speculated that boosting SMIT1 membrane expression would augment receptor-mediated vasodilatation. Adenosine relaxes arteries through Gs coupled A2a/b receptors ^18^ and is far less effective in mesenteric arteries compared to coronary arteries. In control mesenteric arteries adenosine produced a concentration-dependent relaxation that reached approximately 50% at 10 µM, which was prevented by prior application of pan-Kv7 blocker XE991 (Fig 3 A-B). The relaxant effect of adenosine was enhanced after incubation with raffinose for 30 minutes and 4 hours (Fig 3C). In contrast, when XE991 was present, the adenosine-induced relaxation observed under raffinose-treated conditions was significantly reduced (Fig. 3C). Moreover, the relaxant response to ML213 ^15–17^ in mesenteric and coronary arteries was augmented by short term (1h) incubation in raffinose (150 mM; Fig 3D-E). Microelectrode recordings from mesenteric artery smooth muscle cells revealed that incubation with 150 mM raffinose produced a rapid membrane hyperpolarisation (Fig 3H), which was prevented by XE991 (Fig 3I). The Kv7 blocker also induced depolarization of mesenteric vascular smooth muscle cell membrane potential after 60 minutes incubation with raffinose (Fig 3J). Thus, hypertonicity produced a graded and rapidly occurring pro-relaxant effect associated with Kv7 channel-dependent membrane hyperpolarisation.

**Figure 3:**
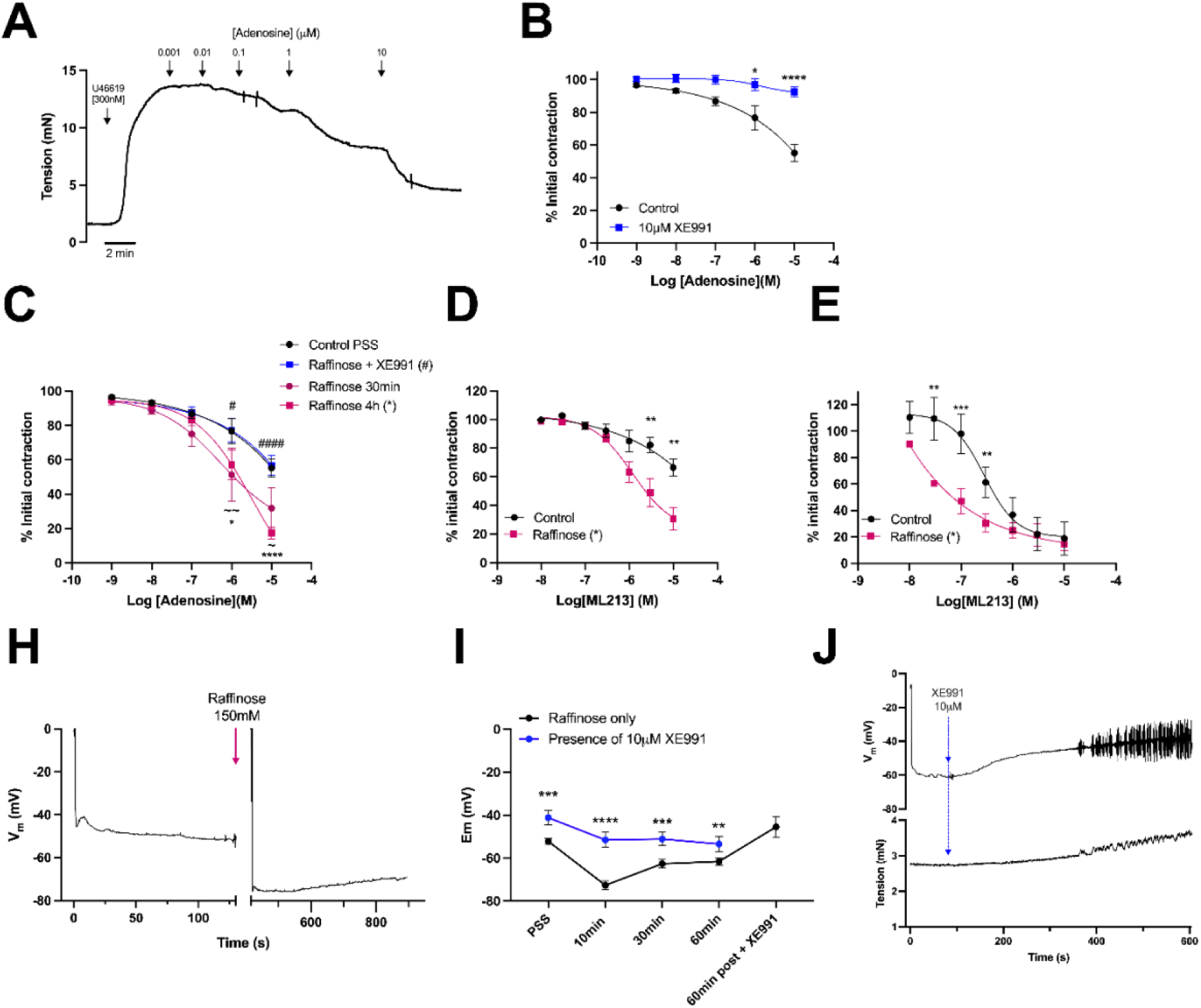
Adenosine and ML213-induced relaxations are enhanced in mesenteric arteries incubated in 150mM raffinose for 2 hours. A representative trace showing the relaxation to Adenosine (1nM - 10µM) in control PSS (A). The mean data of adenosine-induced relaxation in the presence (blue) and absence (black) of 10µM XE991 (B). The mean concentration-dependent relaxation to adenosine (1nM - 10µM) in mesenteric arteries when incubated in either control PSS (black) or 150mM raffinose PSS (pink) for 30 mins (dark pink), 4h (pink) or 150mM raffinose and 10µM XE991 (blue) for 4 hours shown in panel C. The mean response to ML213 in mesenteric arteries (D) and coronary arteries (E) pre-contracted with 300 nM U46619 in the presence (pink, N=6) and absence (black, N=6) of 150mM raffinose. A representative race showing simultaneous measurements of Vm before and after adding 150mM raffinose PSS (H). The mean change in Vm before and after incubating in 150mM raffinose for 10, 30 and 60 minutes in the presence (blue) and absence (black) of XE991 and application of XE991 after incubating in raffinose for 60 minutes (black, I). A representative trace showing the depolarisation upon applying 10 µM XE991 to mesenteric arteries after incubating in 150mM raffinose PSS for 60 minutes is shown in panel J. All data points are shown as mean +-SEM denoted by error bars with N=5, and statistical values are determined by 2-way ANOVA with multiple comparisons are calculated using post hoc Bonferroni test where *=P<0.05, **=P<0.01, ***=P<0.001, ****=P<0.0001.

### 3.3. SMIT1 interacts with Kv7.4/5 channels and Gβγ Subunit to evoke relaxation

Having established the rapid anti-contractile effect of hypertonic external solution experiments were performed to establish the underlying mechanisms. Proximity ligation assay established that in mesenteric artery smooth muscle cels SMIT1 associated with Kv7.4 and Kv7.5 and the interaction was augmented by short term incubation in 150 mM raffinose (Fig 4A-B). Similar data was derived from coronary artery smooth muscle cells (Fig 4C). Gβγ subunits associate with Kv7.4 and enhance channel activity ^19,20^ and this interaction is crucial for the manifestation of several receptor-mediated relaxations ^21^. PLA revealed that Gβγ associated with Kv7.4 and SMIT1 at high levels in mesenteric and coronary arteries and raffinose incubation increased the interaction with Kv7.4 and Gβγ in smooth muscle cells from mesenteric and coronary arteries (figure 4D-F). Thus, extracellular hypertonicity enhanced the association of Kv7 channels with positive regulators SMIT1 and Gβγ.

**Figure 4:**
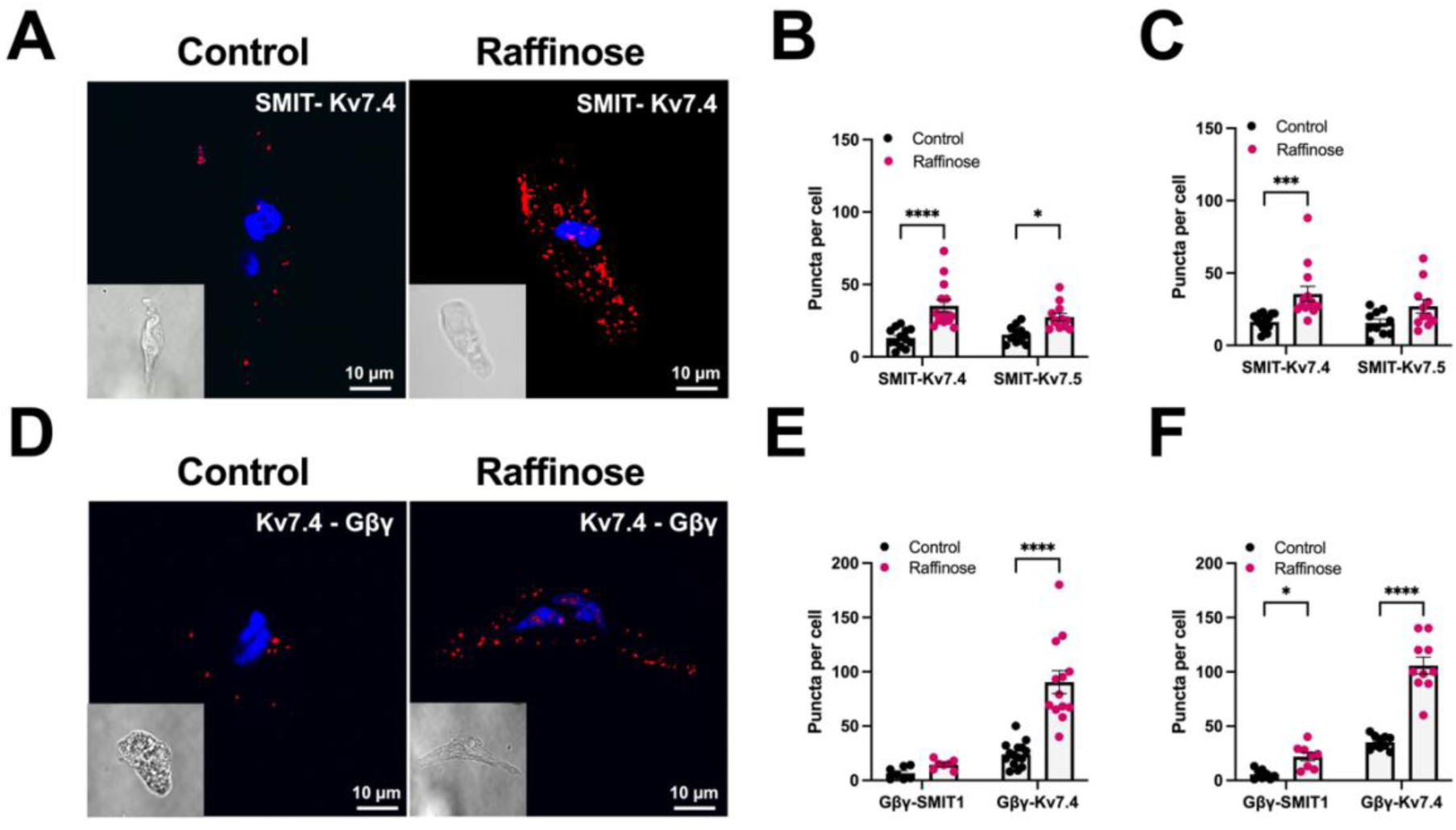
The interaction and role of Kv7 channels and Gβγ in SMIT1 induced relaxations. PLA puncta (red) show the interaction of SMIT with Kv7.4 in mesenteric VSMCs incubated in either control PSS or 150 mM raffinose PSS for 4 hours (A). Bar graph showing the mean number of PLA puncta for SMIT-Kv7.4 and SMIT-Kv7.5 in mesenteric (B) ( N=3 n=15) and coronary (C) (N=3 n=15) VSMCs incubated in either control PSS (black) or 150 mM raffinose PSS (pink) for 4 hours. PLA puncta (red) show the interaction of Kv7.4 with Gβγ in mesenteric VSMCs incubated in either control PSS or 150 mM raffinose PSS for 4 hours (D). Bar chart showing the mean number of Gβγ-Kv7.4 and Gβγ-SMIT1 PLA puncta per cell in mesenteric (E, N=3 n=14) and coronary (F, N=3 n=10) arteries incubated in either control PSS (black) or 150mM raffinose PSS (pink). All data points are shown as mean +-SEM denoted by error bars, and statistical values are determined by 2-way ANOVA with multiple comparisons are calculated using post hoc Bonferroni test where *=P<0.05, **=P<0.01, ***=P<0.001, ****=P<0.0001. (N= number of animals used, n= number of technical repeats).

### 3.4. SMIT1 pro-relaxant effects are mediated by serum glucocorticoid kinase 1 (SGK1)

Incubation in raffinose impaired receptor-mediated contraction within 10 minutes that was not consistent with a hypertonicity-mediated change in SMIT1 gene transcription and subsequent increased protein production. Klaus et al (2008)^10^ showed that cell membrane abundance of SMIT1 was limited by NEDD4.2-mediated ubiquitination, which was down-regulated by serum/ glucocorticoid-dependent kinase 1 (SGK1) phosphorylation. We used proximity ligation assays to show that SGK1 associated with SMIT1, and the interaction was increased upon incubation in 150 mM raffinose in both mesenteric and coronary arteries (Fig 5A-B). Pre-incubation with the SGK1 inhibitor EMD638683 prevented the raffinose-induced increase in SMIT1 membrane abundance (Fig 5C-E). EMD638683 also attenuated the raffinose-induced inhibition of methoxamine in mesenteric (Fig 5F) and coronary arteries (Fig 5G). Rapid alterations of arterial responsiveness by hypertonic media therefore seems to be mediated by recruitment of SGK1 and prompt limiting of SMIT1 recycling.

**Figure 5:**
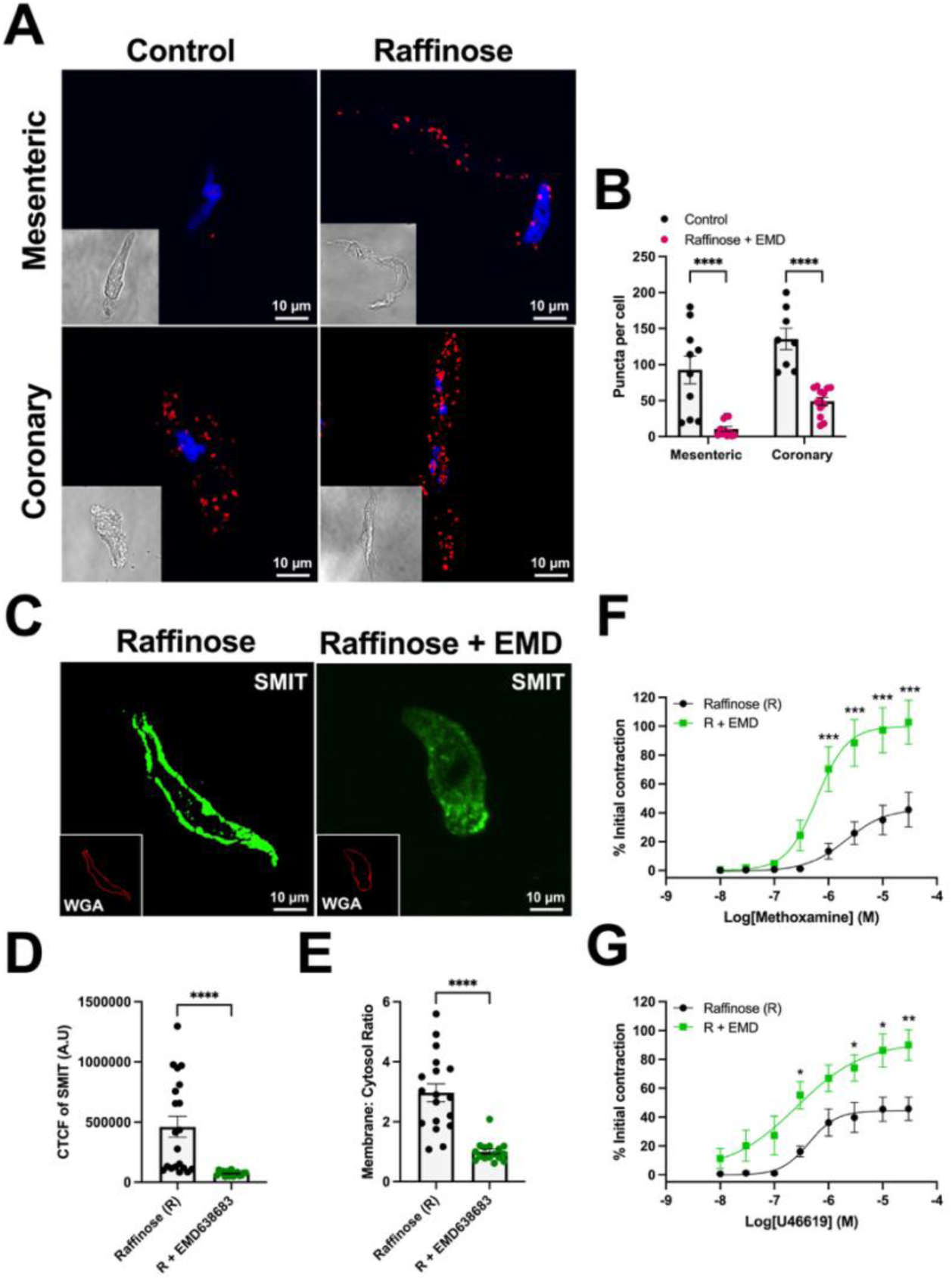
The role of SGK in regulating SMIT1 membrane expression and activity. SMIT-SGK PLA puncta staining (red) in mesenteric arteries and coronary arteries incubated in either control PSS or 150mM raffinose PSS (A). The mean number of PLA puncta in mesenteric (N=3 n=10) and coronary (N=3 n=10) arteries following 4-hour control PSS (black) or 150mM raffinose (pink) incubation in male rats is shown in panel B. Representative staining of SMIT1 (green) in raffinose (C, left) and in raffinose with SGK inhibitor EMD638683 (C, right) with the membrane marker, wheat germ agglutinin (WGA, red) and nucleus stain with DAPI (blue). The mean total cell fluorescence intensity (CTCF) of SMIT1 in mesenteric VSMCs incubated in raffinose alone (black, N=4 n=20) or with an SGK1 inhibitor EMD638683 (green, N=4 n=20) is shown in panel D. The mean membrane: cytosol ratio of SMIT1 staining in mesenteric VSMCs incubated with raffinose alone (black, N=4 n=18) or with EMD638683 (green, N=3 n=16) is shown in panel E. The contraction to methoxamine (10nM - 10µM) following 4-hour incubation in 150mM raffinose in the presence (green) and absence (black) of SGK inhibitor EMD638683 in mesenteric (F, N=7) and coronary (G, N=7) arteries. All data points are shown as mean +-SEM denoted by error bars, and statistical values are determined by 2-way ANOVA with multiple comparisons are calculated using post hoc Bonferroni test where *=P<0.05, **=P<0.01, ***=P<0.001, ****=P<0.0001. (N= number of animals used, n= number of technical repeats).

### 3.5. SMIT1 modulates arterial contractility in female arteries

There is considerable evidence that arterial regulation is sex dependent. Recently, Baldwin et al (2023)^45^ reported that Kv7.2-7.5 activators relaxed several arteries that was influenced by the oestrous cycle stage. We assessed the impact of raffinose in mesenteric and coronary arteries from female rats at different stages of the oestrous cycle segregating them into the higher oestrogen pro-oestrus / oestrus (PE) or lower oestrogen diestrus/ mestestrus (DM) according to Baldwin et al (2023) ^45^. Total SMIT1 abundance was significantly higher in mesenteric arteries from DM females compared to PE females and male rats (Fig 6A-C). In addition, PLA punctae for SMIT1 with Kv7.4, Kv7.5 or Gβγ were higher in smooth muscle cells from DM rats compared to PE females (Fig 6D) and raffinose enhanced the number of associations in cells from DM but not PE rats (Fig 6E-F). A similar pattern of SMIT1 interactions with Kv7 channels and Gβγ was observed in coronary artery smooth muscle cells (Supplement S2-3). Overall, SMIT1 abundance and interaction with Kv7 channels and Gβγ exhibit oestrous cycle dependence.

**Figure 6:**
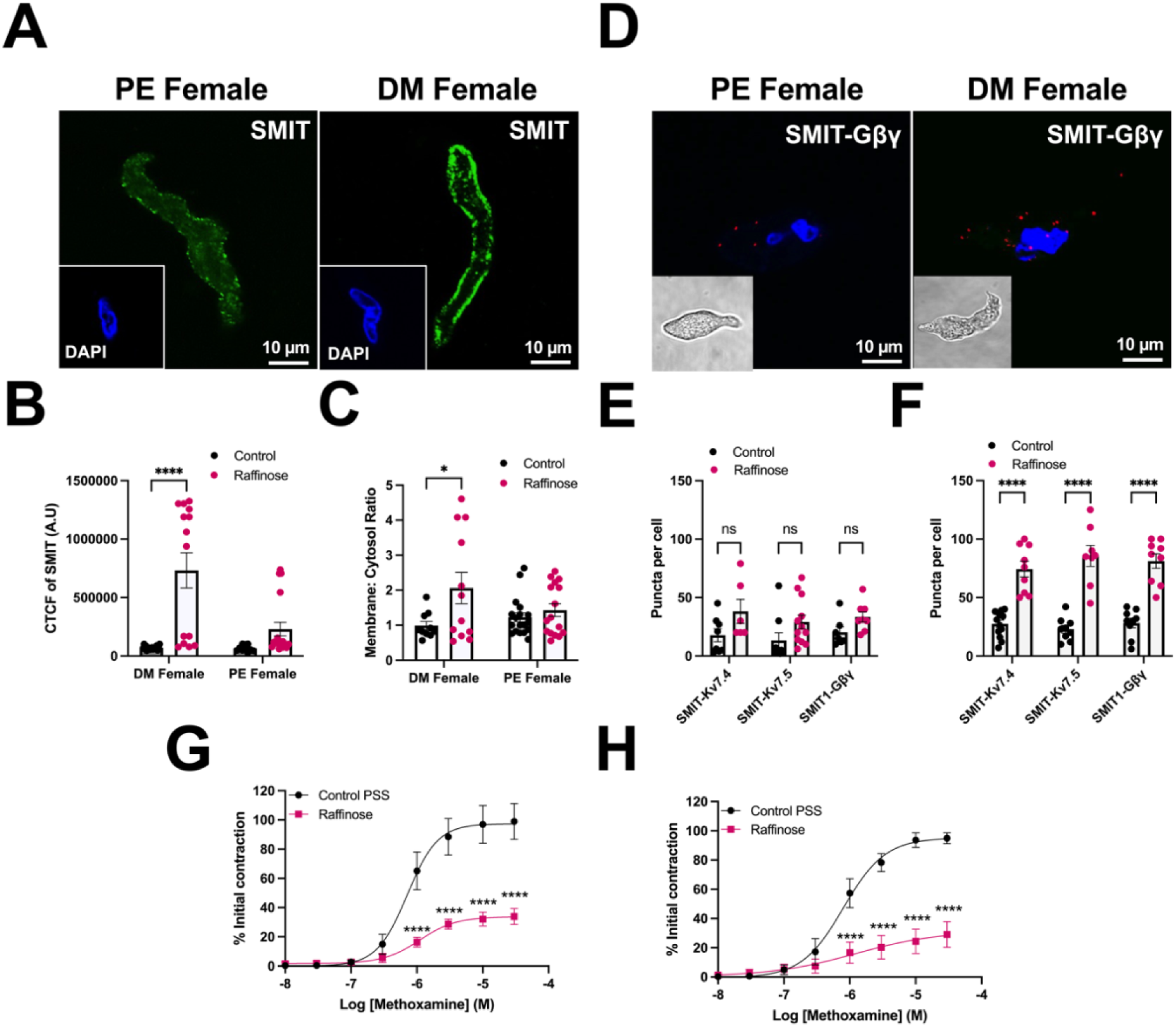
The expression and function of SMIT1 in mesenteric arteries from female rats. A representative image of mesenteric smooth muscle cells stained with SMIT1 (green) and nucleus stain with DAPI in PE female (A, left) and DM female rats (A, left) with and nucleus stain with DAPI (blue). The mean corrected total cell fluorescence (CTCF) intensity of SMIT1 staining in mesenteric arteries from DM female and PE female rats with (pink) and without (black) incubating in 150 mM Raffinose for 4 hours (B). The mean membrane cytosol ratio of SMIT1 in mesenteric arteries from DM female and PE female rats in control vessels (black) or vessels incubated with 150 mM raffinose for 4 hours (pink, C). Proximity ligation assay staining of SMIT-Gβγ puncta in vascular smooth muscle cells from PE and DM female mesenteric arteries with and nucleus stain with DAPI (blue, D). Bar charts showing PLA puncta of SMIT1-Kv7.4, SMIT-Kv7.5 and SMIT-Gβγ interactions in mesenteric arteries from PE female (E, N=2 n=8) and DM female (F, N=3 n=9) rats following incubation for 4 hours in control PSS (black) or 150mM raffinose PSS (pink). The mean concentration-dependent contraction to methoxamine in mesenteric arteries from PE (G) and DM female (H) rats incubated in either control PSS (black) or 150mM raffinose PSS (pink)(N=9). All data points are shown as mean +-SEM denoted by error bars and statistical values are determined by 2-way ANOVA with multiple comparisons are calculated using post hoc Bonferroni test where *=P<0.05, **=P<0.01, ***=P<0.001, ****=P<0.0001.

## 4. Discussion

The interaction between SMIT1 and Kv7 ion channel proteins has been documented in the choroid plexus epithelium, neurones, overexpression systems and more recently arteries demonstrating reciprocal regulation of each other’s activity independent of myo-inositol transport ^2–7^. The present study reveals sodium/ myoinositol transporter SMIT1 to be a crucial regulator of arterial reactivity independent of inositol transport and provides a rapid mechanism for arterial smooth muscle to respond to external stimuli. We show that incubation of arteries in moderately hypertonic solution (150 mM raffinose, 480 mOSm) led to rapid inhibition of receptor-mediated contractions but not high potassium-induced responses in mesenteric arteries. The anti-contractile effect was associated with marked hyperpolarisation of the smooth muscle membrane potential and prevented by molecular knockdown of SMIT1. Conversely, adenosine produced robust relaxations of mesenteric arteries after short term incubation with raffinose that were not present in control arteries and the Kv7 activator ML213 was considerably more potent. Short term incubation of coronary arteries in raffinose also led to impairment of contractile responses to U46619 and methoxamine but augmented relaxations. The membrane induced-hyperpolarisation and pro-relaxant response of raffinose was ameliorated by the Kv7 blocker XE991. Proximity ligation assays revealed that raffinose increased association of SGK1 with SMIT and Gβγ with Kv7 channels. The Gβγ blocker gallein and the SGK1 inhibitor, EMD638683 also suppressed the anti-contractile effect of raffinose. These data reveal that short term increase in extracellular tonicity translates to an anti-contractile phenotype in arteries mediated by SGK1 preventing SMIT1 removal from the cell membrane.

### 4.1. SMIT1 regulates arterial contractility

Overnight incubation (∼16 h) of renal arteries with hypertonic 150mM raffinose and myo-inositol solution has been shown to increase the membrane abundance of SMIT1 and reduced the contractile response to methoxamine and low concentrations of KCl (<40 mM)^6^. The present study advances these observations by showing that incubation in 150 mM raffinose similarly increases SMIT1 expression and reduces methoxamine and U46619 contractions in mesenteric and coronary arteries. However, we discovered that the anti-contractile effect did not require prolonged incubation in hypertonic media as significant inhibition occurred within 30 mins, indicating that SMIT1 can rapidly modulate arterial tone. These findings align with earlier work indicating that acute exposure to hypertonic solutions whether evoked by sucrose, mannitol or high glucose evoked vasorelaxation in several arterial beds including skeletal muscle arterioles, cerebral and coronary arteries through K^+^ channel activation^22–28^.

Acute incubation with raffinose hyperpolarised the resting membrane potential of mesenteric vascular smooth muscle cells, an effect prevented by the Kv7-channel blocker XE991, confirming rapid Kv7 activation under hypertonic conditions with increased SMIT1 expression. Consistent with this, short-term (1 h) exposure to raffinose significantly enhanced relaxations to adenosine and to the Kv7 activator ML213 in both mesenteric and coronary arteries. Conversely, morpholino-based knockdown of SMIT1 protein markedly impaired relaxations to isoprenaline and CGRP, which are mediated partially by recruitment of Kv7 channels ^21,29^, as well as relaxations to ML213. Together these findings indicate SMIT1 is required for full Kv7.4/7.5 channel function and acts as a positive regulator of Kv7-mediated vasodilation, which is consistent with previous studies showing SMIT1 interacts with Kv7.2/3 and Kv7.1 ion channels in neurons and the choroid plexus ^3–5,7^ and the presence of SMIT1 negatively shifts the voltage dependence of these ion channels’ currents and enhances rate of activation ^5^.

Myo-inositol, transported by SMIT1, is a precursor for phosphatidylinositol 4,5-bisphosphate (PIP₂), a key lipid cofactor required for stabilising Kv7 channel opening and promoting activation. Altering SMIT activity has been shown to regulate neuronal excitability by modulating PIP_2_ levels ^3^. Therefore, the inhibitory effects of raffinose in the present study may reflect increased SMIT1-dependent myo-inositol transport and enhanced PIP₂-dependent activation of Kv7.4/7.5 channels. However, no inositol was present in the external solution during the acute experiment or incubation periods. Moreover, the non-selective inhibitor of SMIT, phlorizin, does not prevent the anti-contractile effects produced by raffinose ^6^. Thus, the effects of SMIT1 on Kv7 channels were not due to myo-inositol uptake increasing intracellular PIP_2_ and instead involved direct modulation of Kv7 channels, as indicated by Manville et al 2017 ^5^.

### 4.2. SMIT1 resides close to Kv7.4/5 and Gβγ proteins contributing to vasorelaxation

Co-immunoprecipitation assays showed that SMIT1 interacts with Kv7.2 by binding to the S5-6 pore forming region, altering ion selectivity and channel sensitivity ^5^. In the present study, SMIT1 was found to associate with Kv7.4 and Kv7.5 and this interaction was augmented by short term incubation in 150 mM raffinose in both mesenteric and coronary arteries. Gβγ subunits have been identified as crucial regulators of Kv7.4 ion channel with Gβγ and Kv7.4 interacting in arterial smooth muscle, internal Gβγ enrichment enhancing heterologously expressed Kv7.4 and native Kv7 currrent and various Gβγ blockers including gallein abolishing the voltage-dependent activity of Kv7.4 channels^19^. The present study showed that Gβγ physically interacted with Kv7.4/ 7.5 and SMIT1 in coronary and mesenteric artery smooth muscle cells from male and female rats arteries and these interactions increased following a 4-hour incubation in raffinose-enriched PSS. Moreover, gallein prevented the raffinose-induced reduction in U46619-evoked contractions in mesenteric arteries to a similar extent as the Kv7 inhibitor XE991. As SMIT1 interacts with Gβγ, we propose that the transporter co-ordinates the G-proteins to alter Kv7.4 / 7.5 activity. Hence, Kv7 ion channel proteins, SMIT and Gβγ form a complex to regulate arterial tone. As Kv7 channels in smooth muscle cells is modulated by the single transmembrane auxillary subunit KCNE 4 ^30^ and KCNE proteins modify the effect of SMIT1 on Kv7 channels^4,5^ there is an extra layer of complexity to consider in this signaling complex.

### 4.3. SGK1 regulates SMIT1 expression modulating its effect on arterial contractility

The anticontractile effect of increasing SMIT1 may be due to the raffinose-mediated activation of the transcription factor TonEBP (tonicity-responsive enhancer binding protein), resulting in increased SMIT1 protein expression ^11,31^. However, activation of TonEBP in response to hypertonicity is slow, with a detectible increase only occurring after approximately 3 hours ^11^. The present study observed that the effect of raffinose on contractility manifested rapidly, with U46619- and methoxamine-induced contractions being impaired even after 10 minutes of incubating in high raffinose PSS. Therefore, other regulators of SMIT1 are contributing to its increased membrane protein abundance, resulting in the observed anticontractile effects.

Serum- and glucocorticoid-inducible kinases (SGK1-3) are upregulated by hypertonicity ^32,33^ and are a known post-translational regulator of numerous ion channels ^34–39^ either directly or through regulation of membrane abundance. For instance, SGK modulates the expression of hERG ^37,38^ and NaV1.5 ^34^ proteins in myocardial cell membranes by deactivating the ubiquitin ligase Nedd4-2, thereby preventing the retrograde trafficking of these channels and increasing their membrane abundance. Electrophysiological studies in Xenopus oocytes identified that SMIT1-mediated myo-inositol transport increased when SGK1 was co-expressed due to increased residency of SMIT1 in the membrane ^10^ following phosphorylation and deactivation of Nedd4-2, which removed SMIT1 from the cell membrane. SGK can also phosphorylate SMIT1 *per se*, leading to further SMIT1 membrane expression ^10^. The present study shows that the anti-contractile effect in raffinose incubated arteries was prevented by XE991, gallein and the SGK1 inhibitor, EMD638683. Strikingly, SGK1 was also found to closely interact with SMIT in mesenteric and coronary arteries, with the number of interactions increasing upon raffinose incubation. Our data suggest that the rapid impairment of arterial contractility and pro-relaxant arterial state produced by hypertonicity was caused by upregulated SGK activity enhancing SMIT1 protein expression ^10^ and subsequently increasing SMIT1 interactions with Kv7.4/5 channels. SGK1 also phosphorylates Kv7 channels directly ^40–43^, which could also increase the membrane abundance of Kv7.4/7.5 in the vascular smooth muscle.

### 4.4. Oestrus cycle differences in SMIT1-induced relaxations

The present study demonstrated that the expression of SMIT1 in the presence and absence of 150 mM raffinose was considerably lower in smooth muscle cells from mesenteric and coronary arteries from PE females compared to male and DM females, with DM females consistently having the greatest total fluorescence intensity of SMIT1 protein. Moreover, DM females consistently had a greater number of SMIT1 interactions with Kv7.4, Kv7.5 and Gβγ compared to males and PE females, with PE females overall having considerably fewer SMIT1 interactions in the presence and absence of raffinose. Therefore, there seems to be an oestrus cycle difference in the expression and interactions of SMIT1 in DM female and PE female arteries.Baldwin et al (2022 and 2023)^44, 45^ identified that Kv7 activators were less effective relaxants in arteries from PE females compared to DMfemales due to a reduction in the forward trafficking of Kv7.4 via HSP90 leading to a reduction in Kv7.4 expression in the VSMC membrane ^44,45^. The present work estabdlishes that sex- and oestrous cycledifferences in Kv7 responsiveness may be manifest as a consequence of altered protein-protein interactions in the smooth muscle cell.

### 4.5. Conclusion

This study substantiates the view that SMIT1 is a key component of smooth muscle cell Kv7.4/5 ion channels complexes that regulate arterial contractility and vasodilation in mesenteric and coronary arteries from male and female rats. It coordinates Gβγ interaction with Kv7.4/7.5. Moreover, through a rapid effect involving SGK1 SMIT1 transduces environmental stimuli into arterial responsiveness. These findings highlight a new paradigm in arterial control – the SMIT1-Kv7 chansporter complex.

## Novelty and Significance

Sodium myoinositol transporter 1 or SMIT1 associates with KCNQ-encoded voltage gated potassium channels Kv7.4 and Kv7.5. SMIT1 expression is increased by overnight incubation hypertonic medium, which led to poor responses to different receptor-mediated vasoconstrictors.

*What new information does the article contribute?* Methoxamine- and U46619-mediated contractions in mesenteric resistance and coronary arteries were impaired by short term (30 mins to 4h) incubation in hypertonic bathing solution. Relaxations to adenosine were augmented after 60 mins incubation in hypertonic medium. These pro-relaxant effects were mediated by serum glucocortoid kinase 1 and were underpinned by increased interaction of Kv7 channels with Gβγ subunits.

Novelty and significance. This study identifies the sodium myo-inositol transporter 1 as a key element that transduces environmental stress into change in arterial response through coordination of Kv7 channels with Gβγ proteins.

## Acknowledgments

E.A.F. generated and analysed data, and wrote the manuscript. V.B and A.P.A reviewed the manuscript and contributed to experimental design. C.J.G did the microelectrode recordings. I.A.G. supervised the project, edited the manuscript, and provided the funding for the project. Professor Kim Dora (Oxford University) helped with the analysis of microelectrode experiments.

---

**Graphic Figure:**
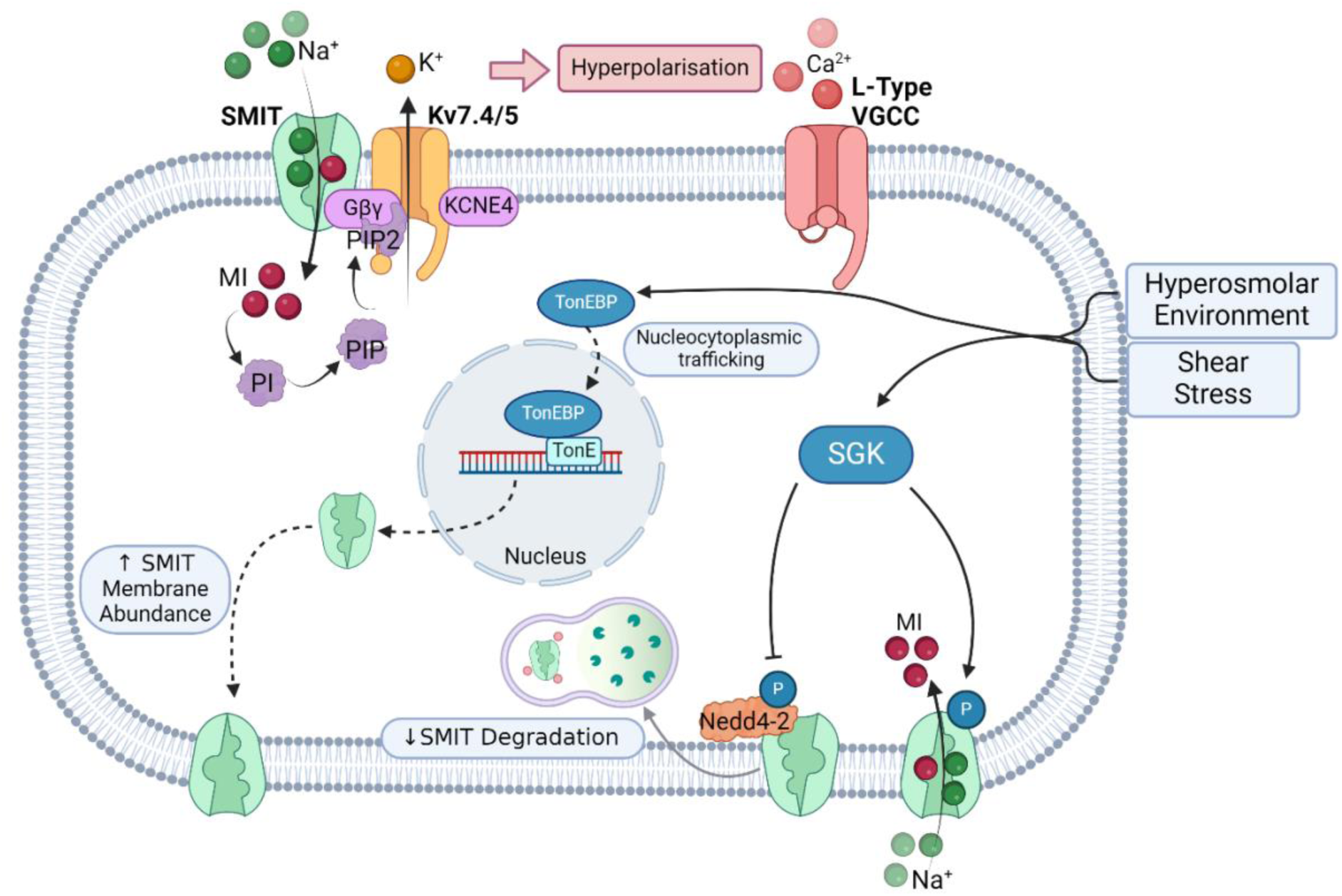
Overview of SMIT1 activity in VSMCs. The tonicity-responsive enhancer binding protein (TonEBP) and serum glucocorticoid kinase (SGK) sense a rise in extracellular hypertonicity and increase SMIT1 membrane abundance in VSMCs. SMIT1 interacts with Kv7.4/5, Gβγ, and KCNE4, promoting hyperpolarisation, VGCC inactivation, and vasorelaxation. (Source: Author using Biorender.com).

